# All about urine: Longitudinal examination of urine pH, specific gravity, proteins, culture, and resistance profiles in healthy dogs

**DOI:** 10.1101/2023.02.28.530482

**Authors:** Andrew McGlynn, Ryan Mrofchak, Rushil Madan, Christopher Madden, Mohammad Jawad Jahid, Dixie Mollenkopf, Thomas Wittum, Sheryl S. Justice, Adam Rudinsky, Jessica Hokamp, Vanessa Hale

## Abstract

**Background:** Urine is routinely evaluated in dogs to assess health. Reference ranges for many urine properties are well established, but the scope of variation in these properties over time within heathy dogs is not well characterized.

**Objectives:** Longitudinally characterize urine properties in healthy dogs over 3 months.

**Animals:** Fourteen healthy client-owned dogs.

**Methods:** Dogs were evaluated for health; then, mid-stream free catch urine was collected from each dog at 12 timepoints: Mornings / afternoons of Days 1, 2, 3; end of Weeks 4, 5, 6, 7 and Months 2 and 3. Urine pH, urine specific gravity (USG), protein, cultures, and antimicrobial resistance profiles were evaluated at each timepoint.

**Results:** Urine pH varied significantly within and between dogs over time (Friedman’s test: within *p* = 0.031; between *p* < 0.005). However, USG, protein, and bacterial richness of urine were consistent within dogs over time, and only varied significantly between dogs (Kruskal-Wallis: between all *p* < 0.005). Antimicrobial resistant isolates were identified in 13 out of 14 dogs with 71% (34 of 48) of the isolates demonstrating resistance to amoxicillin.

**Conclusions and clinical importance:** 1) Urine pH should be assessed at multiple timepoints via pH meter prior to making clinical decisions. 2) Mid-stream free catch urine from multiple healthy dogs yielded high concentrations of bacteria in culture (>10^5^ CFU/mL) and should not be considered the only indicator of urinary tract infection. 3) Bacterial isolates demonstrated widespread resistance to amoxicillin / oxacillin underscoring the need for antimicrobial stewardship.

## Introduction

Urine provides many insights into host health and is routinely included in canine clinical evaluations. Routinely evaluated urine properties include color, pH, urine specific gravity (USG), protein content, and the presence / abundance of chemical compounds such as ketones, bilirubin, and glucose.^1–3^ Urine can also be cultured, and urine sediments evaluated for white blood cells, host cells, and bacteria. Urinalysis aids in screening asymptomatic animals, and provides critical information for diagnostic evaluations of kidney damage, metabolic disease (e.g. diabetes), infection, stone formation or other health conditions.^1,3–5^ While reference intervals for most of these urine properties are well established, the scope and degree of variation within a heathy dog over time are less well defined.

Urine pH, for example, influences urinary tract infection (UTI) and stone formation risk in dogs and can also be monitored to assess response to diets designed to prevent stone formation.^6–8^ Urine specific gravity (USG) tracks concentrating ability of the kidneys and can serve as an indicator of diseases such as kidney disease and diabetes.^1,3,9^ Urine protein profiles also help identify kidney disease, urinary tract inflammation, and distinguish tubular from glomerular damage.^1,10,11^ However, these same urine properties - pH, USG, and urine protein profiles - are also affected by many factors beyond disease, including diet, medications, and hydration status.^1,3,12,13^ Characterizing the normal range of variation in urine properties within a dog over time informs the clinical application and interpretation of these values.

Urine variability can also alter the niches available to commensal bacteria.^14^ Urinary tract commensals are thought to play a role in host health through immune stimulation, colonization resistance, and pathogen clearance.^15–19^ However, few studies have examined the urinary microbiota of dogs via sequencing or culture, and even fewer have evaluated change over time.^15,18,20–24^ Multiple studies have reported that dogs and humans share microbes, including urinary tract pathogens.^25–33^ Thus, characterizing healthy canine urinary microbiota and the associated resistance profiles over time is valuable both for canine health and has implications for One Health.

In this study, we evaluated urine pH, USG, urine protein profiles, and urine culture in 14 healthy dogs over 12 timepoints ranging from a few hours to a few months apart. We also compared pH as measured by dipstick to pH measured via pH meter. Finally, we phenotypically assessed antimicrobial resistance of isolates cultured from the urine of these healthy dogs against amoxicillin, ciprofloxacin, oxacillin, and nalidixic acid. We found that pH was highly variable within dog over time, while urine specific gravity, protein profiles, and microbial culture results were relatively consistent over time. As reported in other studies, the urine dipstick significantly over or underreported urine pH across all pH ranges except for neutral pH 7 ± 0.5. Lastly, we observed resistance against at least one antibiotic in 40 of 48 (83%) tested isolates, representing 12 of 14 healthy dogs (85%).

## Materials and Methods

### Recruiting and Enrollment

All dogs were recruited through the “masked for review” Veterinary Medical Center (IACUC: 2020A00000050). Each dog underwent a physical exam, serum chemistry, complete blood count (CBC), urinalysis, and urine culture prior to enrollment to assess health (**Figure 1**). All dogs were required to be between 1-10 years of age, have a body weight of at least 20 lbs (9 kg), be able to produce ≥ 10 ml of urine in a single urination, have a body condition score of 4 or 5, and be spayed or neutered^34^. Dogs were excluded if they had any history or signs of urinary tract disease, liver or kidney disease, skin infection, gastrointestinal disease, or urogenital abnormalities. Other exclusion criteria included antibiotic use, chemotherapy, or radiation within 3 months of enrollment.

**Figure 1:**
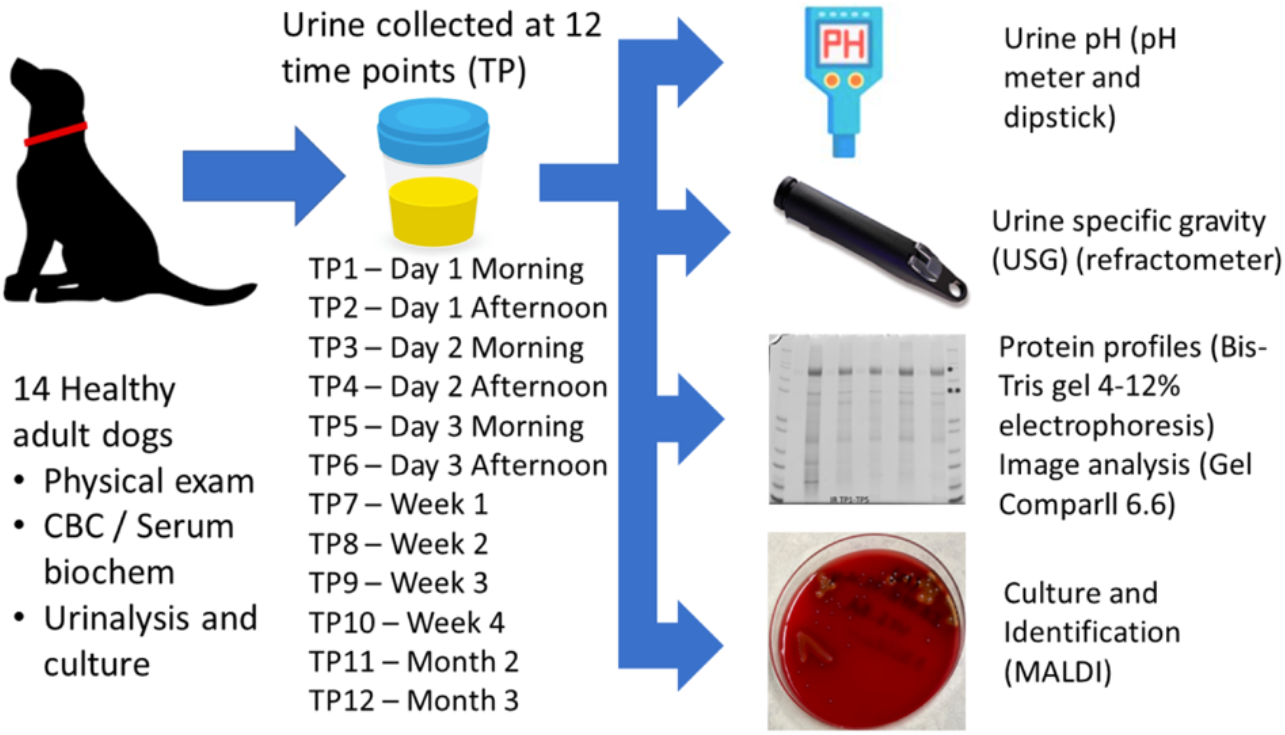
Experimental Design.

### Sample Collection

Mid-stream free catch urine samples were collected from 14 dogs (7 males, 7 females) over 12 timepoints between September 2020 and September 2021 (**Table 1, Supporting Information Supp. Table 1**). The twelve timepoints included: Day 1 Morning (TP1), Day 1 Afternoon (TP2), Day 2 Morning (TP3), Day 2 Afternoon (TP4), Day 3 Morning (TP5), Day 3 Afternoon (TP6), End of Week 1 (TP7), End of Week 2 (TP8), End of Week 3 (TP9), End of Week 4 (TP10), End of Month 2 (TP11), and End of Month 3 (TP12). First-morning urine was collected for all timepoints except the three timepoints that were specifically aimed at collecting “afternoon” urine on Days 1, 2, and 3 (TP2, TP4, TP6). All urine samples were immediately placed on ice following collection and transported to the lab for aliquoting and processing within 6 hours of urination. Urine aliquots designated for pH and USG analysis were brought to room temperature prior to asessment.

**Table 1:** Demographic table. There were no significant differences in age between male and female dog enrolled in this study.

|  | Males | Females | p-value |
| --- | --- | --- | --- |
| Total Number of Dogs | 7 | 7 | 1 |
| Age (mean ± SD) | 3.62 ± 2.55 | 4.50 ± 2.15 | 0.497 |
| Age Range | 1 - 7.08 | 1.25 - 8 | - |

### pH (meter vs. dipstick)

Urine pH was assessed via pH meter (SevenEasy S20, Mettler Toledo, Columbus, OH, United States) and dipstick (Chemstrip^®^ 9, Roche Diagnostics, Rotkreuz, Switzerland). The pH meter was calibrated before measurement using calibration buffer solutions with pH values of 4.00, 7.00, and 10.00. Following calibration, the pH meter probe was submerged in a urine sample until a stable pH reading could be obtained. To measure urine pH via dipstick, one drop (~50 μl) of urine was placed on the dipstick square that evaluates pH. After 1 minute at room temperature, (per manufacturer instructions) the color of the square was matched to a manufacturer guide to assign a pH value to the sample.

### Urine Specific Gravity (USG)

Urine specific gravity was assessed using a refractometer (Reichert VeT 360, Depew, NY, USA). The refractometer was calibrated before usage by placing 1-2 drops (~50 – 100 μl) of deionized, ultrapure water on the refractometer and adjusting to a specific gravity of 1.000 as necessary. Following calibration, 1-2 drops (~50 – 100 μls) of urine was placed on the refractometer and USG was recorded.

### Urine Protein Profiles

Urine protein profiles were generated as described previously in Hokamp et al. 2018 with a few modifications.^10^ In brief, urine was normalized to urine specific gravity (USG) of 0.065 through the addition of ultrapure water. The total combined volume of urine and ultrapure water did not exceed 30 μl, and the water volume was determined based on the following formula: 30 – [0.065/(USG-1)] = μl of ultrapure water added to the urine sample. Ten microliters of lithium dodecyl sulfate (LDS) 4X sample buffer (Bolt™, Thermo Fisher Scientific, Waltham, Massachusetts) was then added to the tube. The tube was briefly vortexed followed by centrifugation for 30 seconds at 400 rcf. Sample tubes were then placed into a heating block for 10 minutes at 70°C. After heating, the samples were vortexed and centrifuged again for 30 seconds at 400 rcf. A gel apparatus (Mini Gel Tank, Thermo Fisher Scientific, Waltham, Massachusetts) was loaded with two 4-12% Bis-Tris gels (Bolt™, Thermo Fisher Scientific, Waltham, Massachusetts) and 2-(N-morpholino)ethanesulfonic acid (MES) running buffer solution (Bolt™, Thermo Fisher Scientific, Waltham, Massachusetts). A standard ladder (Mark12™, Thermo Fisher Scientific, Waltham, Massachusetts) containing polypeptides between 2.5 and 200 kDa, was run in lanes 1 and 12 of each gel. Each urine sample was run in two lanes. The first lane contained the USG-normalized urine samples and in the second lane, referred to as the “MAX loaded” lane, urine was undiluted (not normalized based on USG). Electrophoresis was performed at 200V and 250 mAmps for 30-32 minutes. Gels were subsequently stained with protein stain (Imperial™, Thermo Fisher Scientific, Waltham, Massachusetts) for 2 hours on a low-speed shaker. After staining, gels were placed in ultrapure water on a shaker for 20 minutes to destain. This was repeated 3 times for a total of 3 washes. After the third wash, the gels were placed into ultrapure water on the shaker for an overnight destain. A kimwipe tissue was placed in the corner of the container to absorb excess stain during the destaining process. Gels were then photographed using a biomolecular imager (Amersham™ Typhoon™, GE Healthcare, Chicago, Illinois). Images of each gel were then analyzed and compared based on densitometric curves (GelComparII 6.6, Applied Maths NV, Sint-Martens-Latem, Belgium) generated from protein bands in USG-normalized lanes. MAX-loaded lanes were used for confirmation of band location.

### Urine Culture

Fifty microliters of each urine sample were vortexed then aliquoted into a sterile 2.0 mL microcentrifuge tube. Samples were then vortexed briefly and centrifuged for 1 min. Ten microliters of each urine sample was then plated onto blood agar and MacConkey agar. All plates were then incubated aerobically at 37° C and checked for growth at 24 and 48 hours. Total viable colonies were counted, and all colonies with unique morphologies were picked and individually stored in a 75% glycerol solution at −80°C. Stored samples were later re-plated onto blood agar for 24 hours then subjected to matrix assisted laser desorption ionization-time of flight mass spectrometry (MALDI-TOF) (Bruker Corporation, Billerica, Massachusetts) for bacterial identification. Culture plates with mixed bacterial species were subsequently replated to establish pure cultures, before MALDI-TOF identification.

### Antimicrobial Resistance Profiles

A total of 220 isolates were cultured across all dogs and timepoints. A subset of these isolates (n=48) were then selected for antimicrobial resistance evaluation including: every *Escherichia coli* (n = 4) and *Pseudomonas aeruginosa* (n = 2) isolate based on the association of these bacterial species with UTIs; one isolate from all other bacterial species identified in each dog; and, in some dogs from which we cultured the same bacterial species repeatedly, we selected the first and last isolate of that bacterial species cultured per dog.^35^ The selected isolates were then grown on the blood agar plates and incubated for 18-24 hours at 37°C. Colonies from each isolate were then picked and inoculated into 3 ml of double distilled water and normalized to a 0.5 McFarland standard at 625 nanometers with turbidity range of 0.08-0.1. The normalized bacterial solution was then streaked on Muller Hinton Agar (MHA) supplemented with antimicrobials at breakpoint concentrations reported in dogs and/or specific to the canine urinary tract. Gram-negative isolates were cultured on 3 different MHA plates: one containing nalidixic acid (32 ug/mL), one containing ciprofloxacin (4 ug/mL), and one containing amoxicillin (8 ug/mL).^35–38^ Gram-positive isolates were cultured on 2 different MHA plates: one containing oxacillin (0.5 ug/mL), and one containing amoxicillin (8 ug/mL).^39^ All plates were incubated for 18-24 hours at 37°C then checked for bacterial growth. Amoxicillin, a beta-lactam, was selected because it is a first line antimicrobial for lower UTIs.^40^ Oxacillin was selected to evaluate resistance against a second beta-lactam antibiotic. Ciprofloxacin, a fluoroquinolone, was selected because fluoroquinolones are the recommended alternatives to beta-lactams for UTIs in the case of beta-lactam resistance.^40^ Nalidixic acid, a first-generation quinolone, was selected to parse potential quinolone versus fluoroquinolone resistance.

### Statistical analyses

All analyses were performed in R Studio version 4.1.0 and statistical significance was assessed at a *p*-value of 0.05. Data on urine pH, specific gravity, and protein profiles were tested for normality using a Shapiro-Wilks Test. To test for differences by dog and by time, we employed Kruskal-Wallis, Pairwise Wilcoxon Rank Sum, and Friedman’s tests.

## Results

### pH within and between dogs over time

Urine pH was highly variable both within and between dogs. The average pH across all dogs and all timepoints as measured by pH meter was 6.54 ± 0.83 (range: 5.32 – 8.93), and as measured by dipstick was 6.35 ± 1.15 (range: 5.0 – 9.0). The largest pH ranges recorded in single dogs over time (12 time points) were: 5.45 – 8.31 by pH meter in dog HF and 5 – 9 by dipstick in three dogs –ArB, FC, HF. The smallest pH ranges recorded in single dogs over time were: 5.56-6.98 by pH meter in dog AbB, and by dipstick, 5 – 7 in three dogs (AbB, IR, LM) and 7 – 9 in one dog (GC). pH values from both meter and dipstick were not normally distributed (Shapiro-Wilk Normality Test, *p* = 3.809 x 10^-6^ and 1.614 x 10^-10^, respectively) and varied significantly between dogs (pH meter: Kruskal-Wallis, *p* = 5.021 x 10^-5^; dipstick: Kruskal-Wallis, *p* = 0.0002, **Figure 2A, B**) and over time within dogs (pH meter: Friedman’s, *p* = 0.031; dipstick: Friedman’s, *p* = 0.007, **Figure 2A, B**). Dog GC had a significantly higher pH as measured by meter and dipstick than almost all other dogs (all Wilcoxon pairwise, *p* < 0.05; **Supporting Information Supp. Tables 2, 3**). There was no significant difference in pH between males and females (meter: Kruskal-Wallis, *p* = 0.075, dipstick: Kruskal-Wallis, *p* = 0.151).

**Figure 2:**
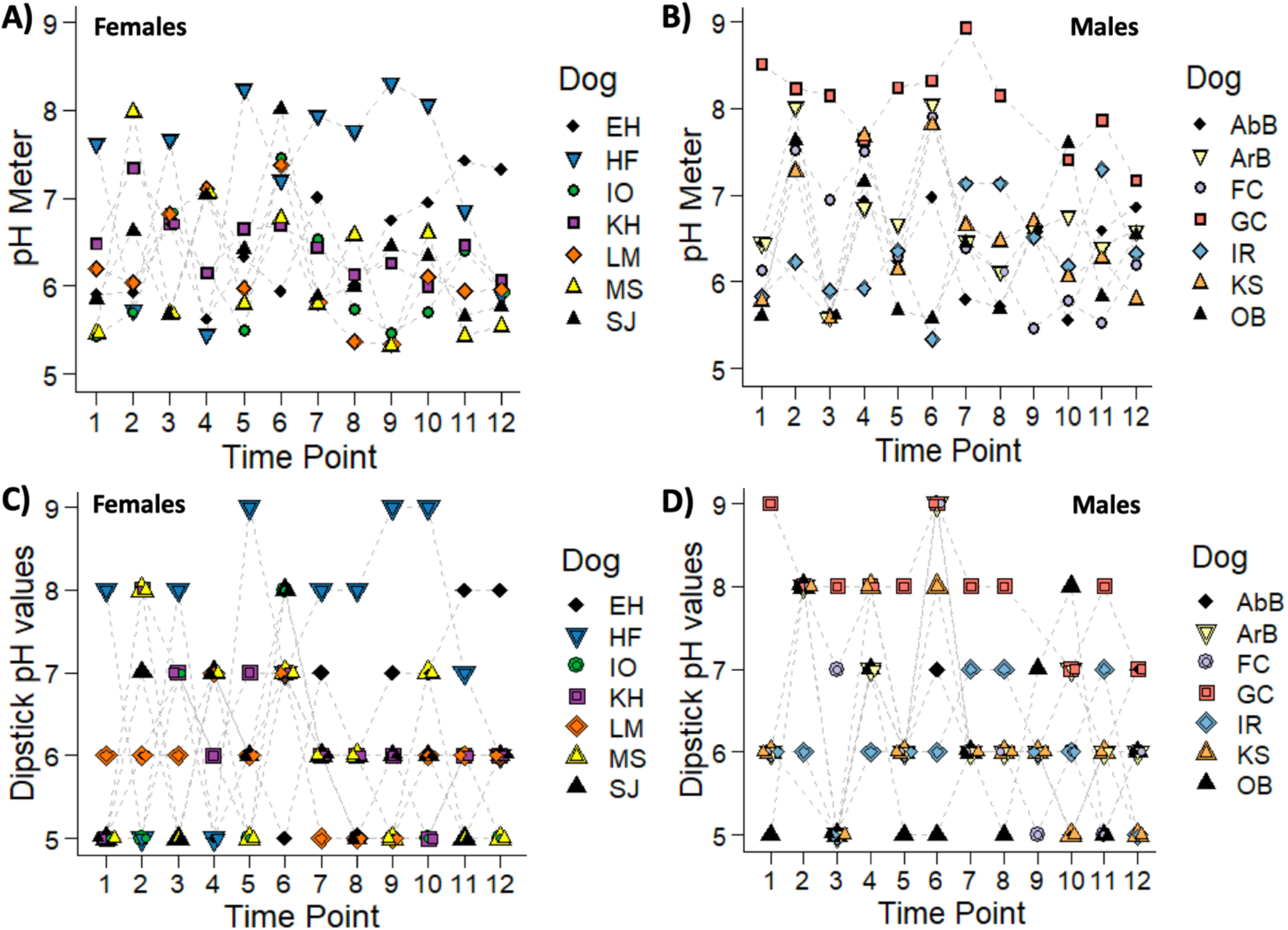
Urine pH of dogs over 12 timepoints. Urine pH as measured by pH meter in a) female dogs and b) male dogs. pH by meter varied significantly between dogs (Kruskal-Wallis, *p* = 5.021 x 10^-5^) and within dogs over time (Friedman’s test, *p* = 0.031). Urine pH as measured by dipstick in c) female dogs and d) male dogs. pH by dipstick varied significantly between dogs (Kruskal-Wallis, *p* = 0.0002) and within dogs (Friedman’s test, *p* = 0.007). There was no significant difference in pH between males and females by both methods (meter: Kruskal-Wallis, *p* = 0.075, dipstick: Kruskal-Wallis, *p* = 0.151).

### pH meter versus dipstick

Overall, urine pH measured via meter differed significantly from pH measured via dipstick (Kruskal-Wallis, *p* = 0.042). The mean difference (pH meter – dipstick pH = difference) across all samples was 0.187 (**Figure 3**) while the mean absolute difference across all samples was 0.419. When we compared differences in pH readings between pH meter and dipsticks (pH meter – dipstick pH = difference), we found that the mean difference across all samples was 0.187 (**Figure 3A**). Above pH 7.5, dipsticks are less accurate than the pH meter, consistently overestimated pH at basic pHs. When pH was acidic (pH < 6.5), differences indicated that dipsticks consistently underestimated pH. To determine if specific pH ranges resulted in greater differences between pH meter and dipstick readings, we grouped samples into four categories based on pH meter: pH <5.5 (n = 10 samples), pH 5.5 – 6.49 (n = 81 samples), pH 6.50 – 7.49 (n = 47 samples), and pH ≥7.5 (n = 27 samples). We then compared the absolute value of differences between pH measurement methods across these four groups. The lowest difference (most similarity) between pH meter and dipstick values was in the neutral range group (pH 6.50 – 7.49) (mean difference ± SD = 0.350 ± 0.231). At pH values below 6.5 and above 7.5, the absolute differences between methods were greater (**Figure 3B**); although, average differences did not differ significantly between groups (Kruskal-Wallis, *p* = 0.32).

**Figure 3:**
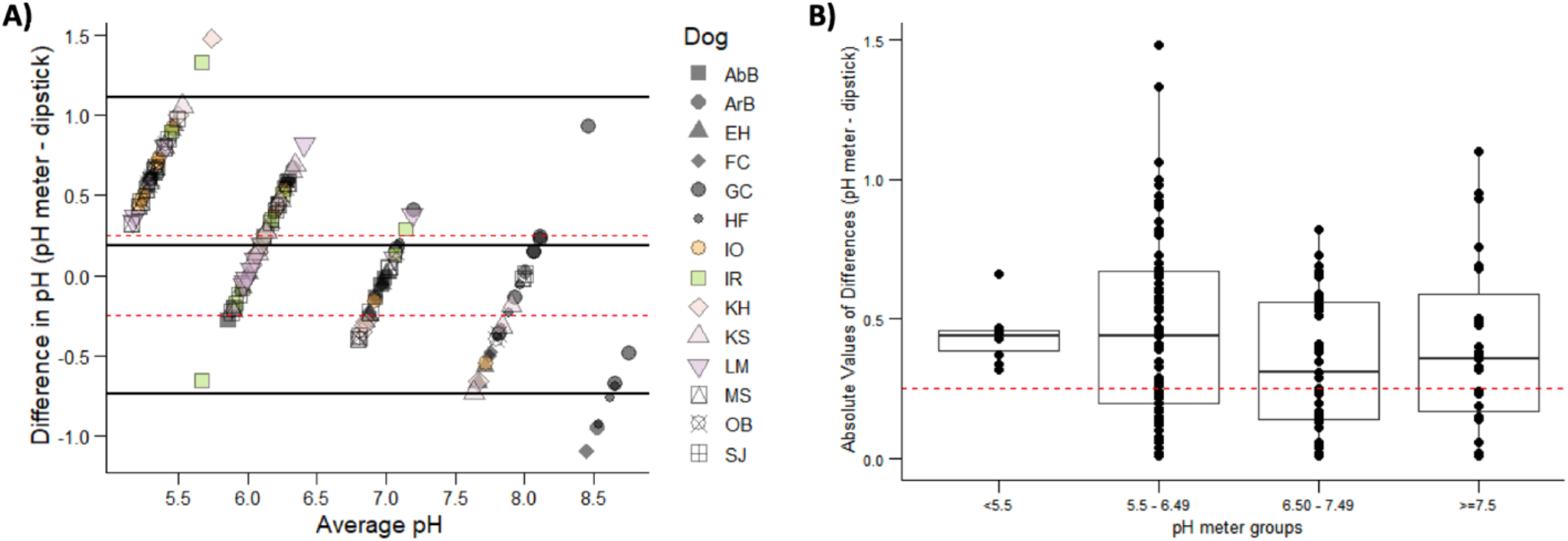
Comparison of urine pH as measured by pH meter vs dipstick. **a**) Bland-Altman plot showing calculated differences in pH (pH meter pH – dipstick pH). Bold black lines represent mean difference (0.187) and 95% CI interval. Dashed red lines represent the clinically significant threshold for differences between methods (0.25 and −0.25). At acidic pHs, the dipstick consistently underestimated pH while at basic pHs the dipstick consistently overestimated pH. **b**) Box-and-whiskers plot showing absolute differences in pH by group: pH <5.5, pH 5.5-6.49, pH 6.50-7.49, pH ≥7.5. There were no significant differences between groups (Kruskal-Wallis, *p* = 0.3204) although differences were higher above and below the neutral pH range (6.5-7.49). Dashed red line indicates the clinically significant threshold for differences between methods (0.25).

### USG differences between morning and afternoon

Healthy dogs generally exhibited limited variation in USG over time. The average USG across all dogs and all timepoints was 1.040 ± 0.011 (range: 1.010 – 1.060). USG values were not normally distributed (Shapiro-Wilk Normality Test, *p* = 0.0003) and varied significantly between dogs (Kruskal-Wallis, *p* = 9.099 x 10^-16^, **Figure 4A, B**), but not within dogs over time (Friedman’s, *p* = 0.377, **Figure 4A, B**). The average difference between minimum and maximum USG within dogs was 0.021 ± 0.009 (range: 0.009-0.038). Males had a significantly higher USG values compared to females (Kruskal-Wallis, *p* = 0.002). In pairwise comparisons, dogs LM, IO, and MS (all females) had significantly lower USGs than most other dogs (Wilcoxon pairwise, most *p* < 0.05, **Supporting Information Supp. Table 4)**. To evaluate if USG values varied significantly between first-morning and afternoon urine, we analyzed a subset of samples from timepoints 1 – 6 for all 14 dogs. Morning USG values were higher (mean ± SD = 1.043 ± 0.009) than afternoon USG values (1.038 ± 0.011); although, this difference was not statistically significant (Friedman’s, *p* = 0.1655, **Figure 4C**), likely due to a relatively small sample size. These results suggest that within-day variation in USG was not as strong as the variation observed between different dogs.

**Figure 4:**
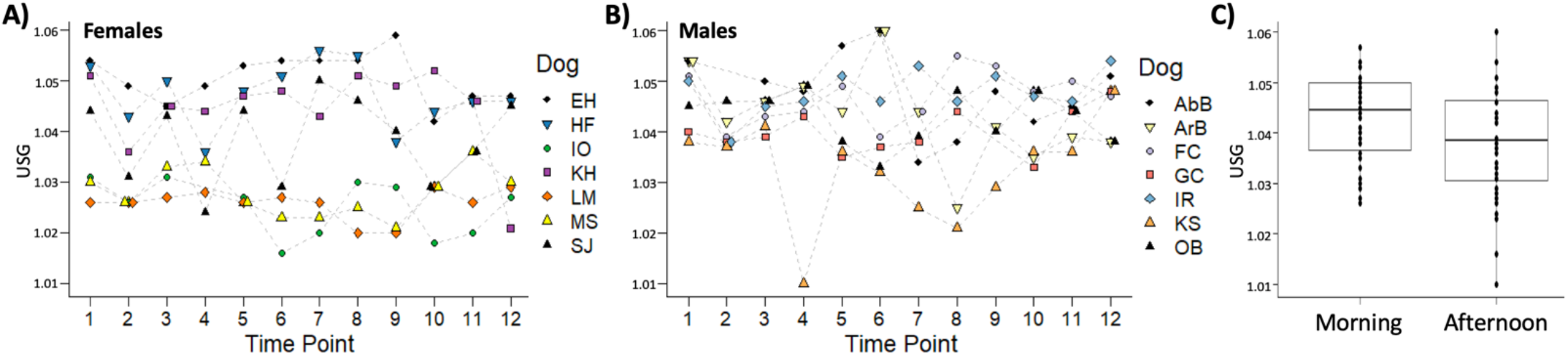
Urine specific gravity (USG) of dogs over time. USG of **a**) female dogs and **b**) male dogs over 12 timepoints. USG, measured via refractometer, varied significantly between dogs (Kruskal-Wallis, *p* = 9.099 x 10^-16^) but not within dog over time (Friedman’s test *p* = 0.377). Males had a significantly higher USG values compared to females (Kruskal-Wallis, p = 0.002). c) USG values compared between first-morning urine and afternoon urine. Morning USG were higher than afternoon USG, all this difference was not significant (Friedman’s test, *p* = 0.166).

### Urine protein profiles

We evaluated total band number and relative surface area of protein bands in the 4-12% Bis-Tris gels. Total band number estimates protein richness or the number of different types of proteins present while relative surface area is a proxy for protein concentration. Total band number ranged from 0-6 across all samples, and differed significantly between dogs (Kruskal-Wallis, test, *p* = 2.105 x 10^-9^) and by sex (Kruskal-Wallis, *p* = 0.056) but not within dogs over time (Friedman test, *p* = 0.191) (**Figure 5A, B**; **Supporting Information Supp. Table 5)**. The two most commonly detected protein bands displayed apparent molecular weights consistent with albumin (54.86-58.95 kDa) and Tamm-Horsfall protein (82.74-86.34 kDa)^10,41^. A few protein bands with apparent molecular weights between 9.10 and 47.31 kDa were also observed occasionally. The nature of these proteins is currently unknown, and they occurred infrequently and at low concentrations compared to albumin and Tamm-Horsfall. The relative surface area of protein bands consistent with albumin and Tamm-Horsfall differed significantly between dogs (Kruskal-Wallis test: albumin *p* = < 2.2 x 10^-16^; Tamm-Horsfall *p* = < 2.2 x 10^-16^**); Supporting Information Supp. Tables 6-7**), but did not differ significantly within dogs over time (Friedman’s: albumin *p* = 0.346; Tamm-Horsfall *p* = 0.606). Albumin concentration did not differ significantly by sex (Kruskal-Wallis, *p* = 0.535, **Figure 5C**), but Tamm-Horsfall concentration was significantly higher in males compared to females (Kruskal-Wallis, *p* = 0.045, **Figure 5D**). There was no significant correlation found between pH and total protein band number, albumin concentration or Tamm-Horsfall concentration (total protein band number: *R* = 0.057, *p* = 0.36; albumin: *R* = 0.067, *p* = 0.22; Tamm-Horsfall: *R* = −0.0099, *p* = 0.86; **Supporting Information Supp. Figure 1**).

**Figure 5:**
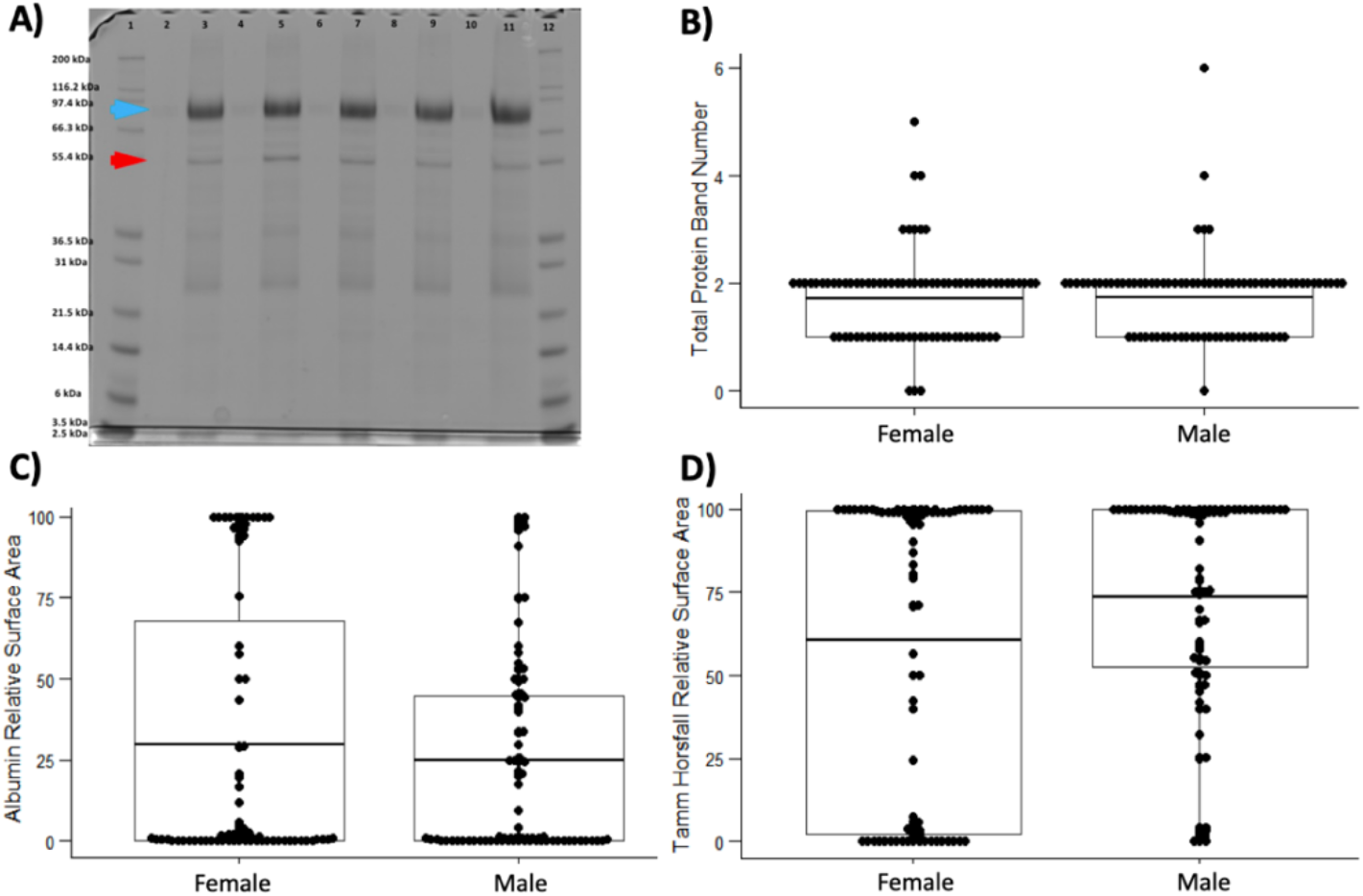
Urine protein profiles. **a)** An example gel from dog KH. Lanes are numbered and ladders are included in lanes 1 and 12. Lanes 2 and 3 contain urine samples from TP 6, Lanes 4 and 5 are samples from TP 7, Lanes 6 and 7 are samples from TP 8, Lanes 8 and 9 are samples from TP 9, and Lanes 10 and 11 are samples from TP 10. Lanes 2, 4, 6, 8, 10 contain USG-normalized urine while lanes 3, 5, 7, 9, 11 are “MAX Loaded” loaded lanes that contain urine not normalized by USG. All densitometric evaluations were performed on USG-normalized lanes. MAX-loaded lanes were used for confirmation of band location. Red arrow denotes protein bands ~58.95-54.86 kD, which is consistent with albumin. Blue arrow denotes bands around ~86.34-82.74 kDa, which is consistent with Tamm-Horsfall protein. Tamm-Horsfall and albumin bands were the most commonly observed bands in the urine protein profiles. **b)** Total band number and **c)** albumin concentration did not differ significantly by sex (Kruskal-Wallis: total band number *p* = 0.056; albumin *p* = 0.535). **d)** Tamm-Horsfall concentration was significantly higher in males compared to females (Kruskal-Wallis, *p* = 0.048).

### Urine cultures

Bacteria was cultured in 50% (85/168) of the urine samples collected over 12 timepoints in 14 dogs. The most commonly cultured bacteria were *Streptococcus canis* and *Staphylococcus pseudintermedius* (**Figure 6A**). Three out of 14 healthy dogs (IR, KS, HF) exhibited urine cultures with > 10^5^ CFU/mL, and two of these dogs (IR, HF) cultured > 10^5^ CFU/mL at more than one timepoint. In all but one case, these cultures were exclusively composed of *S. canis* or *S. pseudintermedius*. In dog HF, one time point (TP11) included a mixed culture of *<u>S. pseudintermedius</u>, E. coli*, and *Bacillus marisflvai*. There was no significant difference in the number of colonies observed at 24 or 48 hours on blood agar or MacConkey agar (Kruskal-Wallis: blood agar *p* = 0.503; MacConkey *p* = 0.771; **Supporting Information Supp. Table 8**). The presence or absence of aerobic culturable bacteria did not differ by sex (Fisher exact test, *p* = 1). Furthermore, the number of total bacterial taxa (richness) cultured aerobically in each dog did not differ significantly by sex (t-test, p = 0.53, **Figure 6B**) or timepoint (Friedman test, p= 0.32), but did differ significantly by dog (Friedman test, p= 4.1 x 10^-10^). Specifically, dog OB’s cultures exhibited significantly greater bacterial diversity than most other dogs in this study, and *Citrobacter* spp. were consistently cultured at 11/12 (91.7%) timepoints. The eight most common taxa cultured at multiple timepoints and in multiple dogs were: *Streptococcus canis, Staphylococcus pseudintermedius, Curtobacterium flaccumfaciens, Pantoea agglomerans, Haemophilus haemoglobinophilus, Escherichia coli, Lysinibacillus fusiformes*, and *Staphylococcus intermedius* (**Supporting Information Supp. Table 9**). However, most taxa were cultured intermittently at fewer than 5 timepoints, and not found as consistently as Staphylococcus or *Streptococcus* spp., except for *Citrobacter* spp. in dog OB.

**Figure 6:**
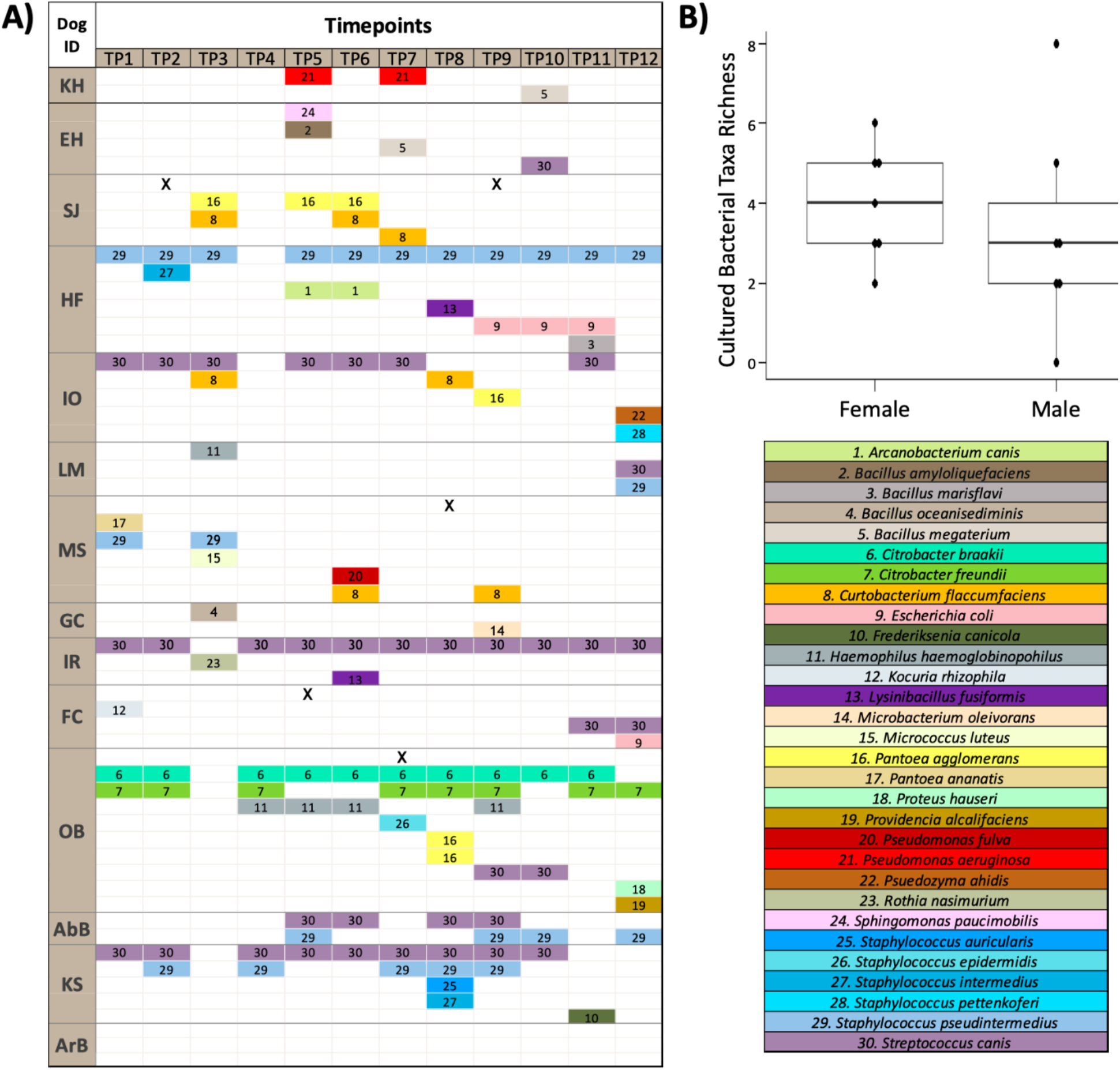
Urine bacterial culture results. **a**) Bacterial species cultured by dog and over time. **b)** Bacterial taxa richness by sex. Number of total bacterial taxa types (richness) aerobically cultured in each dog did not differ significantly by sex (t-test, p = 0.53) or timepoint (Friedman test, p= 0.32, Figure 3) but did differ significantly by dog (Friedman test, p= 4.1 e-10). X indicates that an organism was cultured but not able to be identified by MALDI-TOF. *Per MALDI: Citrobacter species are challenging to speciate and these taxonomic assignments are probable but not definite.

### Antimicrobial resistance profiles

We characterized the resistance profiles of 48 bacterial isolates derived from the urine cultures described above. This included 17 gram-negative isolates and 31 gram-positive isolates (**Supporting Information Supp. Table 10**). The 17 gram-negative isolates were cultured from 7 dogs (4 females, 3 males). OB grew the greatest number of unique gram-negative taxa (n=6). All 17 (100%) gram-negative isolates were resistant to amoxicillin (at 8 ug/mL), and 3 of 17 (17.6%) were resistant to nalidixic acid (at 32 ug/mL). None of the gram-negative taxa were resistant to ciprofloxacin (at 4 ug/mL). The 31 gram-positive isolates were cultured from 13 dogs (6 males, 7 females). Seventeen out of 31 (54.8%) of the gram-positive isolates were resistant to amoxicillin (at 8 ug/mL), while 14 out of 31 (45.2%) were resistant to oxacillin (0.5 ug/mL). Eight gram-positive isolates were resistant to both oxacillin and amoxicillin while 15 isolates were resistant to either oxacillin or amoxicillin but not both. Seven gram-positive isolates were susceptible to both oxacillin and amoxicillin. In a few cases, the same taxa from the same dog had differing resistance profiles over time. For example, in dog AbB, a *Staphylocuccus pseudintermedius* at timepoint 5 (Day 3 Morning) displayed resistance to amoxicillin, but at timepoint 12 (End of Month 3), *S. pseudintermedius* from AbB was not resistant to amoxicillin. Differing taxa within the same dog at the same timepoint also displayed differing resistance profiles. For example, in dog KS at timepoint 8, a *Staphylococcus intermedius* demonstrated resistance to both oxacillin and amoxicillin while *S. auricularis* present in the same dog at the same timepoint, was only resistant to amoxicillin.

## Discussion

In this study, we longitudinally evaluated urine pH, specific gravity, protein, and culture and antimicrobial resistance profiles in 14 healthy dogs over a three-month period. Urine pH varied significantly within and between dogs over time. However, USG, urine protein, and the number of taxa cultured from urine were consistent within dogs over time, and only varied significantly between dogs. Only one dog, OB, consistently cultured bacterial species other than *Staphylococcus* and *Streptococcus* spp. suggesting a urinary bacterial signature unique to this individual dog. However, the scope of this study was to analyze longitudinal bacteria by routine clinical urinalysis culture media. Further research using Enhanced Qualitative Urinary Culture (EQUC) may provide insight into the longevity of viable urobiome taxa by culture-based methods.^42^ Evidence for antimicrobial resistance was identified in 13 out of 14 healthy dogs with the majority of isolates (34 of 48, 71%) demonstrating resistance to amoxicillin.

Urine pH was highly variable within and between dogs. This was not unexpected as urine pH is influenced by multiple factors including diet, disease, age of urine specimen, drug therapies, and bacterial types present in the urine/bladder.^43^ While we did not control for diet in this study, all dogs were confirmed to be healthy based on physical exam, blood work, and urinalysis, and no dog reported any urinary tract issues or infections in the 18 months following enrollment in this study (**Supporting Information Supp. Table 11**). We also processed all urine samples within 6 hours of urination, limiting the potential for pH changes due to specimen handling. Eight dogs (26 total samples accounting for multiple timepoints) exhibited a urine pH outside of, and specifically higher than, the healthy canine urine pH range (5.0 – 7.5) established by Chew et al. 2011.^1^ This indicates that healthy canine urine can vary outside this range. Urine from 11 dogs also exhibited variation from acidic to basic - ranging from pH at or below 6 to pH at or above 7.25, indicating that urine pH was not consistently acidic or consistently basic in most dogs. In comparing pH meter versus dipstick, pH values differed significantly, with the meter producing higher values on average than the dipsticks; although, dipsticks tended to overestimate pH at basic pH values. Additionally, the average absolute difference in pH between meter and dipstick was 0.419 (range 0.01-1.48), which exceeds a previously established clinically acceptable difference of 0.25 between pH measurement methods. ^5,44^ Our results support previous findings indicating a poor concordance between pH meter and pH dipstick values.^6,44,45^ Due to the dynamic nature of urine pH observed in healthy canines, pH readings at multiple timepoints via pH meter are recommended prior to making clinical decisions (including differential diagnoses and treatment) involving urine pH management.

The USG values observed in this study (1.01 – 1.060) fall within a previously established canine USG reference range (1.010-1.070).^46^ USG values were also relatively consistent within a dog over time, with a mean difference of 0.021 ± 0.009, similar to a previously study that reported a mean difference of 0.015 ± 0.007 within dogs over a week.^47^ Although USG can fluctuate based on factors like hydration status, our results indicate that a single USG measurement from a healthy dog will generally be representative of that dog’s USG.^47^ Unlike van Vonderen et al. 1997, we did not observe a significant difference in USG between first morning and afternoon urine; however, average USG was lower in the afternoon in our study and our sample size (n=14 dogs) was small compared to the van Vonderen et al. 1997 study (n=89 dogs) suggesting that temporal differences in USG are small but consistent.^13^ As such, first morning urine is still recommended for USG measurements. We also found that USG was significantly higher in males as compared to females. Higher USG in males has been reported in other species but not dogs.^48–51^ Notably, males in this study were, on average, but not significantly, younger than females (mean ± SD: males 3.6 ± 2.5; females 4.5 ± 2.1). A previous study reported a 0.001 unit decline in canine USG for each increasing year of age; thus, age may be driving the sex difference observed in USG here.^47^

Like USG, urine protein profiles differed significantly between dogs but not within dogs over time. The two most commonly detected protein bands were consistent with Tamm-Horsfall protein and albumin. Tamm-Horsfall, a tubular protein involved in immune defense, and small amounts of albumin (≤30 mg/dL), a plasma protein, are considered normal findings in canine urine.^1,52–54^ Differences in protein profiles between dogs may be driven by age or breed. Increased protein loss through urine is typically observed as dogs age and renal filtration function declines through irreversible nephron loss or glomerulosclerosis, which is more common in older animals.^55^ The number of nephrons per kidney may also vary widely between breeds, impacting renal filtration.^55^ While gel electrophoresis has been used in previous studies to investigate canine urine protein profiles in diseases like chronic kidney disease, pyometra, and leptospirosis, this is the first study, to our knowledge, that characterizes urine protein profiles in healthy dogs over time.^10,56,57^

Urine culture results were, unsurprisingly, dominated by skin-associated microbes (*Staphylococcus* and *Streptococcus spp*.), as urine was collected midstream free-catch.^58^ In general, cystocentesis or catheterization is recommended for culturing to avoid skin and genital contaminants. In addition to skin microbes, we also observed several potential urinary tract pathogens in culture including *E. coli* and *P. aeruginosa*. However, these taxa, at low abundances, can be part of the of the normal urinary tract microbiota in dogs, and the presence of these taxa in asymptomatic individuals does not warrant treatment.^15,23,24,59^ Notably, three dogs cultured high concentrations bacteria (>10^5^ CFU/mL) at least once. This bacterial concentration in midstream free-catch urine (10^5^ CFU/mL) has been suggested as clinically significant for UTI, but the bacteria we observed at these concentrations were generally skin commensals (*S. pseudintermedias, S. canis).^60^* Additionally, all of these dogs remained healthy, asymptomatic, and never exhibited pyuria during and after the study (**Supporting Information Supp. Table 11**). These results demonstrate the potential for high-level contamination in healthy free catch urine, as has been noted previously.^35^

In relation to antibiotic resistance profiles, 93% of the dogs (13 out of 14) and 83.3% of the isolates demonstrated resistance to beta-lactams (amoxicillin, oxacillin or both). Only three gram negative isolates – all *Psuedomonas spp* - displayed resistance to quinolones (nalidixic acid), and no gram-negative isolates demonstrated resistance to fluoroquinolones (ciprofloxacin). *Pseudomonas* species are commonly resistant to quinolones and fluoroquinolones^61,62^. The absence of resistance to fluoroquinolones was considered positive as these drugs are typically reserved for beta-lactam resistant infections.^35,40^ However, the overwhelming resistance to beta-lactams was concerning considering the source: Healthy dogs that had not received any antibiotics in at least 3 months, with most not having received antibiotics for over a year or more. Other studies have reported similarly wide-spread resistance to beta-lactams in bacterial isolates from dogs, including isolates from healthy dogs.^63–65^ Amoxicillin is one of the most commonly used antibiotics in veterinary medicine and is the most frequently prescribed antibiotic for canine UTIs in the United States.^66,67^ The abundant presence of amoxicillin-resistant bacteria isolated in the urine of healthy dogs raises several questions: Is resistance being promoted through prior exposure to a beta-lactam given frequency of use? Is resistance being acquired or transferred from the dog’s environment (e.g., soil, water, diet) or from other hosts (e.g., other pets or humans within a household) as it has already been established that dogs and humans can share urinary tract microbes and pathogens?^25–33^ Given that amoxicillin / beta-lactams are also used to treat a variety of human infections, what are the public health implications of high resistance burdens in healthy dogs that share our households?

## Conclusions

This study is a comprehensive examination of healthy canine urine including pH, USG, protein, culture, and resistance profiles. This work lays the foundation for longitudinal characterization of urine profiles within and between healthy dogs over time. Key take-aways from this study on healthy dogs include: 1) Urine pH varied widely over time indicating that pH should be assessed at more than one timepoint via pH meter prior to making clinical decisions based on pH. 2) USG and protein results were relatively stable over time, suggesting that measurement of these properties at a single timepoint can portray an accurate representation of that dog’s true values. 3) Mid-stream free catch urine from multiple healthy dogs yielded high concentrations of bacteria in culture (>10^5^ CFU/mL) confirming that free catch urine can be highly contaminated and such concentrations of skin bacteria in asymptomatic dogs should not be considered an indicator of a UTI. 4) Urine bacterial isolates demonstrated widespread resistance to amoxicillin and oxacillin underscoring the critical need for antimicrobial stewardship in practice.

## Abbreviations used in the manuscript

CBC: complete blood count
CFU: colony-forming unit
IACUC: Institutional Animal Care and Use Committee
LDS: lithium dodecyl sulfate
MALDI-TOF: matrix assisted laser desorption ionization-time of flight mass spectrometry MES - 2-(N-morpholino)ethanesulfonic acid
MHA: Muller Hinton Agar
TP: timepoint
USG: urine specific gravity
UTI: urinary tract infection

## Acknowledgements

We would like to thank the following: The Ohio State University College of Veterinary Medicine Canine Funds for the financial support of the project, The Muffin Sniadowski Student Research Endowment Fund for providing student financial support to AM for summer research, The Undergraduate Research Apprentice Program for providing financial support to RM, the clients and dogs involved in this study, and Dr. Hannah Klein for the help on physical exams.

## IACUC

IACUC #2020A00000050 at the Ohio State University College of Veterinary Medicine

