## Supporting Information for "All about urine: Longitudinal examination of urine pH, specific gravity, proteins, culture, and resistance profiles in healthy dogs"

### Supplemental Tables

**Supplemental Table 1:** Age, sex, and breed of all enrolled dogs. MN = neutered male, FS = spayed female.

| ID | Sex | Breed | Age (in years) |
|---|---|---|---|
| EH | FS | Spaniel Mix | 8 |
| HF | FS | Borzoi | 6 |
| KH | FS | Mixed Breed | 5 |
| SJ | FS | Pit Mix (American Staffordshire Terrier) | 4.92 |
| IO | FS | Dalmatian | 3 |
| MS | FS | Mixed Coonhound | 3 |
| LM | FS | Dalmatian | 1.6 |
| OB | MN | Doberman Pinscher | 7.08 |
| AbB | MN | Border Collie Mix | 6 |
| ArB | MN | Australian Shepherd | 5 |
| GC | MN | Cardigan Welsh Corgi | 4 |
| IR | MN | Gordon Setter | 1.25 |
| FC | MN | Great Dane/English Mastiff Mix | 1 |
| KS | MN | Golden/Lab Mix | 1 |
| **Average Age of Dogs Enrolled** | | | **4.06 ± 2.31** |
| **Age Range** | | | **1-8 years** |

**Supplemental Table 2: Pairwise comparisons of pH by pH meter between dogs.** GC had a significantly higher average pH by meter than almost all other dogs (Wilcox pairwise, most $p < 0.05$). *denotes statistical significance

| | AbB | ArB | EH | FC | GC | HF | IO | IR | KH | KS | LM | MS | OB |
|---|---|---|---|---|---|---|---|---|---|---|---|---|---|
| ArB | 1 | - | - | - | - | - | - | - | - | - | - | - | - |
| EH | 1 | 1 | - | - | - | - | - | - | - | - | - | - | - |
| FC | 1 | 1 | 1 | - | - | - | - | - | - | - | - | - | - |
| GC | 0.0074* | 0.0363* | 0.0109* | 0.0364* | - | - | - | - | - | - | - | - | - |
| HF | 1 | 1 | 1 | 1 | 1 | - | - | - | - | - | - | - | - |
| IO | 1 | 1 | 1 | 1 | 0.0126* | 0.7373 | - | - | - | - | - | - | - |
| IR | 1 | 1 | 1 | 1 | 0.0065* | 1 | 1 | - | - | - | - | - | - |
| KH | 1 | 1 | 1 | 1 | 0.0065* | 1 | 1 | 1 | - | - | - | - | - |
| KS | 1 | 1 | 1 | 1 | 0.0288* | 1 | 1 | 1 | 1 | - | - | - | - |
| LM | 1 | 1 | 1 | 1 | 0.0065* | 1 | 1 | 1 | 1 | 1 | - | - | - |
| MS | 1 | 1 | 1 | 1 | 0.0139* | 1 | 1 | 1 | 1 | 1 | 1 | - | - |
| OB | 1 | 1 | 1 | 1 | 0.0139* | 1 | 1 | 1 | 1 | 1 | 1 | 1 | - |
| SJ | 1 | 1 | 1 | 1 | 0.0139* | 1 | 1 | 1 | 1 | 1 | 1 | 1 | 1 |

**Supplemental Table 3: Pairwise comparisons of pH by dipstick between dogs.** GC had a significantly higher average pH by dipstick than almost all other dogs (Wilcox pairwise, most $p < 0.05$). *denotes statistical significance

| | AbB | ArB | EH | FC | GC | HF | IO | IR | KH | KS | LM | MS | OB |
|---|---|---|---|---|---|---|---|---|---|---|---|---|---|
| ArB | 1 | - | - | - | - | - | - | - | - | - | - | - | - |
| EH | 1 | 1 | - | - | - | - | - | - | - | - | - | - | - |
| FC | 1 | 1 | 1 | - | - | - | - | - | - | - | - | - | - |
| GC | 0.01* | 0.1926 | 0.1079 | 0.4755 | - | - | - | - | - | - | - | - | - |
| HF | 1 | 1 | 1 | 1 | 1 | - | - | - | - | - | - | - | - |
| IO | 1 | 1 | 1 | 1 | 0.0223* | 0.5994 | - | - | - | - | - | - | - |
| IR | 1 | 1 | 1 | 1 | 0.0063* | 1 | 1 | - | - | - | - | - | - |
| KH | 1 | 1 | 1 | 1 | 0.0251* | 1 | 1 | 1 | - | - | - | - | - |
| KS | 1 | 1 | 1 | 1 | 0.1331 | 1 | 1 | 1 | 1 | - | - | - | - |
| LM | 1 | 1 | 1 | 1 | 0.0049* | 0.9671 | 1 | 1 | 1 | 1 | - | - | - |
| MS | 1 | 1 | 1 | 1 | 0.0251* | 1 | 1 | 1 | 1 | 1 | 1 | - | - |
| OB | 1 | 1 | 1 | 1 | 0.0676 | 1 | 1 | 1 | 1 | 1 | 1 | 1 | - |
| SJ | 1 | 1 | 1 | 1 | 0.0208* | 1 | 1 | 1 | 1 | 1 | 1 | 1 | 1 |

**Supplemental Table 4: Wilcox pairwise comparisons of urine specific gravity (USG) between dogs.** LM, IO, and MS had significantly lower average USGs than most other dogs (Wilcox pairwise, most $p < 0.05$). *denotes statistical significance

| | AbB | ArB | EH | FC | GC | HF | IO | IR | KH | KS | LM | MS | OB |
|---|---|---|---|---|---|---|---|---|---|---|---|---|---|
| ArB | 1 | - | - | - | - | - | - | - | - | - | - | - | - |
| EH | 1 | 1 | - | - | - | - | - | - | - | - | - | - | - |
| FC | 1 | 1 | 1 | - | - | - | - | - | - | - | - | - | - |
| GC | 1 | 1 | 0.0333* | 0.698 | - | - | - | - | - | - | - | - | - |
| HF | 1 | 1 | 1 | 1 | 1 | - | - | - | - | - | - | - | - |
| IO | 0.0073* | 0.0285* | 0.0047* | 0.0049* | 0.0049* | 0.005* | - | - | - | - | - | - | - |
| IR | 1 | 1 | 1 | 1 | 0.2351 | 1 | 0.0048* | - | - | - | - | - | - |
| KH | 1 | 1 | 1 | 1 | 1 | 1 | 0.0285* | 1 | - | - | - | - | - |
| KS | 0.0981 | 0.8419 | 0.0084* | 0.0245* | 1 | 0.0578 | 1 | 0.0239* | 0.3451 | - | - | - | - |
| LM | 0.0046* | 0.0326* | 0.0029* | 0.003* | 0.003* | 0.003* | 1 | 0.0029* | 0.0326* | 1 | - | - | - |
| MS | 0.0073* | 0.0309* | 0.0031* | 0.0033* | 0.0077* | 0.0037* | 1 | 0.0032* | 0.053* | 1 | 1 | - | - |
| OB | 1 | 1 | 0.2341 | 1 | 1 | 1 | 0.0049* | 1 | 1 | 0.3146 | 0.003* | 0.006* | - |
| SJ | 1 | 1 | 0.0575 | 1 | 1 | 1 | 0.1665 | 0.1417 | 1 | 1 | 0.0491* | 0.4997 | 1 |

**Supplemental Table 5: Pairwise comparisons of total protein band number between dogs.** SJ had a significantly greater number of bands than four other dogs while FC and KS had significantly fewer bands than four other dogs (Wilcox pairwise, most $p < 0.05$). *denotes statistical significance

| | AbB | ArB | EH | FC | GC | HF | IO | IR | KH | KS | LM | MS | OB |
|---|---|---|---|---|---|---|---|---|---|---|---|---|---|
| ArB | 1 | - | - | - | - | - | - | - | - | - | - | - | - |
| EH | 1 | 1 | - | - | - | - | - | - | - | - | - | - | - |
| FC | 1 | 0.1417 | 0.4704 | - | - | - | - | - | - | - | - | - | - |
| GC | 0.3735 | 1 | 1 | 0.0053* | - | - | - | - | - | - | - | - | - |
| HF | 1 | 1 | 1 | 1 | 0.2147 | - | - | - | - | - | - | - | - |
| IO | 1 | 1 | 1 | 1 | 1 | 1 | - | - | - | - | - | - | - |
| IR | 1 | 1 | 1 | 0.1181 | 1 | 1 | 1 | - | - | - | - | - | - |
| KH | 1 | 1 | 1 | 1 | 0.072 | 1 | 1 | 0.7803 | - | - | - | - | - |
| KS | 1 | 0.4704 | 1 | 1 | 0.0248* | 1 | 1 | 0.3035 | 1 | - | - | - | - |
| LM | 1 | 1 | 1 | 1 | 0.072 | 1 | 1 | 0.7803 | 1 | 1 | - | - | - |
| MS | 0.2741 | 1 | 1 | 0.0068* | 1 | 0.1551 | 1 | 1 | 0.0618 | 0.0253* | 0.0618 | - | - |
| OB | 0.2741 | 1 | 1 | 0.0068* | 1 | 0.1551 | 1 | 1 | 0.0618 | 0.0253* | 0.0618 | 1 | - |
| SJ | 0.1611 | 1 | 1 | 0.0068* | 1 | 0.0913 | 0.6325 | 1 | 0.0425* | 0.0205* | 0.0425* | 1 | 1 |

**Supplemental Table 6: Pairwise comparisons of albumin concentration between dogs.** Most dogs differed significantly in relative surface areas of albumin, which is a proxy for protein concentrations. For example, GC, LM, and ArB had significantly increased albumin concentrations compared to most other dogs (Wilcox pairwise, most $p < 0.05$). *denotes statistical significance

| | AbB | ArB | EH | FC | GC | HF | IO | IR | KH | KS | LM | MS | OB |
|---|---|---|---|---|---|---|---|---|---|---|---|---|---|
| ArB | 0.0067* | - | - | - | - | - | - | - | - | - | - | - | - |
| EH | 1 | 0.0139* | - | - | - | - | - | - | - | - | - | - | - |
| FC | 1 | 0.0128* | 1 | - | - | - | - | - | - | - | - | - | - |
| GC | 0.004* | 1 | 0.0033* | 0.0017* | - | - | - | - | - | - | - | - | - |
| HF | 1 | 0.0024* | 1 | 1 | 0.0024* | - | - | - | - | - | - | - | - |
| IO | 1 | 1 | 1 | 1 | 1 | 1 | - | - | - | - | - | - | - |
| IR | 1 | 0.011* | 1 | 1 | 0.0033* | 1 | 1 | - | - | - | - | - | - |
| KH | 1 | 0.0024* | 1 | 1 | 0.0024* | 1 | 1 | 1 | - | - | - | - | - |
| KS | 1 | 0.0655 | 1 | 1 | 0.0034* | 1 | 1 | 1 | 1 | - | - | - | - |
| LM | 0.0027* | 0.11 | 0.0023* | 0.0011* | 0.3278 | 0.0016* | 0.2737 | 0.0023* | 0.0016* | 0.0014* | - | - | - |
| MS | 0.0052* | 1 | 0.0033* | 0.0101* | 1 | 0.0024* | 1 | 0.0087* | 0.0024* | 0.0339* | 0.081 | - | - |
| OB | 0.004* | 1 | 0.0033* | 0.0037* | 0.0183* | 0.0024* | 1 | 0.0069* | 0.0024* | 0.0591 | 0.004* | 1 | - |
| SJ | 0.0145* | 1 | 0.0054* | 0.0646 | 1 | 0.0024* | 1 | 0.0221* | 0.0024* | 0.0907 | 0.0907 | 1 | 1 |

**Supplemental Table 7: Pairwise comparisons of Tamm Horsfall concentration between dogs.** Most dogs differed significantly in relative surface areas of Tamm-Horsfall protein, which is a proxy for protein concentrations. For example, GC, LM, and ArB had significantly decreased Tamm-Horsfall concentrations compared to most other dogs (Wilcox pairwise, most $p < 0.05$). *denotes statistical significance

| | AbB | ArB | EH | FC | GC | HF | IO | IR | KH | KS | LM | MS | OB |
|---|---|---|---|---|---|---|---|---|---|---|---|---|---|
| ArB | 0.0067* | - | - | - | - | - | - | - | - | - | - | - | - |
| EH | 1 | 0.0139* | - | - | - | - | - | - | - | - | - | - | - |
| FC | 1 | 0.0128* | 1 | - | - | - | - | - | - | - | - | - | - |
| GC | 0.004* | 1 | 0.0033* | 0.0017* | - | - | - | - | - | - | - | - | - |
| HF | 1 | 0.0024* | 1 | 1 | 0.0024* | - | - | - | - | - | - | - | - |
| IO | 0.003* | 0.1012 | 0.0029* | 0.0012* | 0.0625 | 0.0019* | - | - | - | - | - | - | - |
| IR | 1 | 0.0221* | 1 | 1 | 0.0033* | 1 | 0.0029* | - | - | - | - | - | - |
| KH | 1 | 0.0024* | 1 | 1 | 0.0024* | 1 | 0.0019* | 1 | - | - | - | - | - |
| KS | 1 | 0.6146 | 1 | 1 | 0.0655 | 1 | 0.0109* | 1 | 1 | - | - | - | - |
| LM | 0.0027* | 0.11 | 0.0023* | 0.0011* | 0.3278 | 0.0016* | 1 | 0.0023* | 0.0016* | 0.0112* | - | - | - |
| MS | 0.004* | 1 | 0.0033* | 0.0101* | 1 | 0.0024* | 0.0127* | 0.0087* | 0.0024* | 0.4317 | 0.081 | - | - |
| OB | 0.004* | 1 | 0.0033* | 0.0028* | 0.0781 | 0.0024* | 0.0067* | 0.0054* | 0.0024* | 0.4739 | 0.0074* | 1 | - |
| SJ | 0.0112* | 1 | 0.0087* | 0.0413* | 1 | 0.0024* | 0.1427 | 0.0345* | 0.0024* | 0.8006 | 0.4024 | 1 | 1 |

**Supplemental Table 8: Number of colonies present at 24 and 48 hours.** Urine (10 µl) from each dog was cultured aerobically on blood agar and MacConkey agar at all timepoints. Colonies were counted at 24 and 48 hours. There was no significant difference in the number of colonies present at 24 or 48 hours (Kruskal-Wallis: blood agar $p = 0.503$; MacConkey $p = 0.771$). Four out of 14 healthy dogs (OB, IR, KS, HF) exhibited urine cultures with > $10^5$ CFU/mL, and two of these dogs (OB, IR), both male, cultured > $10^5$ CFU/mL at >5 timepoints. NA= no sample obtained or data recorded at this timepoint. TNTC = too numerous to count.

| | | Total Colony Counts | | | |
|---|---|---|---|---|---|
| **Dog** | **TP** | **Blood Agar 24 hours** | **Blood Agar 48 hours** | **MaConkey Agar 24 hours** | **MaConkey Agar 48 hours** |
| FC | 1 | 0 | 1 | 0 | 0 |
| FC | 2 | 0 | 0 | 0 | 0 |
| FC | 3 | 0 | 0 | 0 | 0 |
| FC | 4 | 0 | 0 | 0 | 0 |
| FC | 5 | 0 | 0 | 0 | 1 |
| FC | 6 | 0 | 0 | 0 | 0 |
| FC | 7 | 0 | 0 | 0 | 0 |
| FC | 8 | 0 | 0 | 0 | 0 |
| FC | 9 | 0 | 0 | 0 | 0 |
| FC | 10 | 0 | 0 | 0 | 0 |
| FC | 11 | 3 | 3 | 0 | 0 |
| FC | 12 | 13 | 10 | 0 | 0 |
| AbB | 1 | 0 | 0 | 0 | 0 |
| AbB | 2 | 0 | 0 | 0 | 0 |
| AbB | 3 | 0 | 0 | 0 | 0 |
| AbB | 4 | 0 | 0 | 0 | 0 |
| AbB | 5 | 4 | 4 | 0 | 0 |
| AbB | 6 | 2 | 3 | 0 | 0 |
| AbB | 7 | 0 | 0 | 0 | 0 |
| AbB | 8 | 1 | 1 | 0 | 0 |
| AbB | 9 | 4 | 4 | 0 | 0 |
| AbB | 10 | 7 | 6 | 0 | 0 |
| AbB | 11 | 0 | 0 | 0 | 0 |
| AbB | 12 | 1 | 2 | 0 | 0 |
| ArB | 1 | 0 | 0 | 0 | 0 |
| ArB | 2 | 0 | 0 | 0 | 0 |
| ArB | 3 | 0 | 0 | 0 | 0 |
| ArB | 4 | 0 | 0 | 0 | 0 |
| ArB | 5 | 0 | 0 | 0 | 0 |
| ArB | 6 | 0 | 0 | 0 | 0 |
| ArB | 7 | 0 | 0 | 0 | 0 |
| ArB | 8 | 0 | 0 | 0 | 0 |
| ArB | 9 | 0 | 0 | 0 | 0 |
| ArB | 10 | 0 | 0 | 0 | 0 |
| ArB | 11 | 0 | 0 | 0 | 0 |
| ArB | 12 | 0 | 0 | 0 | 0 |
| EH | 1 | 0 | 0 | 0 | 0 |
| EH | 2 | 0 | 0 | 0 | 0 |
| EH | 3 | 0 | 0 | 0 | 0 |
| EH | 4 | 0 | 0 | 0 | 0 |
| EH | 5 | 2 | 2 | 0 | 0 |
| EH | 6 | 0 | 0 | 0 | 0 |
| EH | 7 | 1 | 1 | 0 | 0 |
| EH | 8 | 0 | 0 | 0 | 0 |
| EH | 9 | 0 | 0 | 0 | 0 |
| EH | 10 | 0 | 1 | 0 | 0 |
| EH | 11 | 0 | 0 | 0 | 0 |
| EH | 12 | 0 | 0 | 0 | 0 |
| GC | 1 | 0 | 0 | 0 | 0 |
| GC | 2 | 0 | 0 | 0 | 0 |
| GC | 3 | 1 | 1 | 0 | 0 |
| GC | 4 | 0 | 0 | 0 | 0 |
| GC | 5 | 0 | 0 | 0 | 0 |
| GC | 6 | 0 | 0 | 0 | 0 |
| GC | 7 | 0 | 1 | 0 | 0 |
| GC | 8 | 0 | 0 | 0 | 0 |
| GC | 9 | 0 | 7 | 0 | 0 |
| GC | 10 | 0 | 0 | 0 | 0 |
| GC | 11 | 0 | 0 | 0 | 0 |
| GC | 12 | 0 | 0 | 0 | 0 |

### Supplemental Table 8 continued

| | | Total Colony Counts | | | |
|---|---|---|---|---|---|
| Dog | TP | Blood Agar 24 hours | Blood Agar 48 hours | MaConkey Agar 24 hours | MaConkey Agar 48 hours |
| HF | 1 | 46 | 46 | 0 | 0 |
| HF | 2 | 6 | 6 | 0 | 0 |
| HF | 3 | TNTC | TNTC | 0 | 0 |
| HF | 4 | 0 | 0 | 0 | 0 |
| HF | 5 | 3 | 0 | 0 | 0 |
| HF | 6 | 8 | 12 | 0 | 0 |
| HF | 7 | 12 | 15 | 0 | 0 |
| HF | 8 | 2 | 5 | 0 | 0 |
| HF | 9 | 10 | 1 | 0 | 0 |
| HF | 10 | 8 | 21 | 0 | 0 |
| HF | 11 | TNTC | TNTC | 2 | 2 |
| HF | 12 | 14 | 14 | 0 | 0 |
| IO | 1 | 13 | 15 | 0 | 0 |
| IO | 2 | 2 | 2 | 0 | 0 |
| IO | 3 | 3 | 6 | 0 | 0 |
| IO | 4 | NA | NA | NA | NA |
| IO | 5 | 6 | 5 | 0 | 0 |
| IO | 6 | 2 | 3 | 0 | 0 |
| IO | 7 | 0 | 15 | 0 | 3 |
| IO | 8 | 1 | 1 | 0 | 0 |
| IO | 9 | 0 | 0 | 2 | 2 |
| IO | 10 | 0 | 0 | 0 | 0 |
| IO | 11 | 2 | 1 | 0 | 0 |
| IO | 12 | 7 | 7 | 0 | 0 |
| IR | 1 | ~1000 | ~1000 | 0 | 0 |
| IR | 2 | ~1000 | ~1000 | 0 | 0 |
| IR | 3 | 56 | 52 | 0 | 1 |
| IR | 4 | 84 | 86 | 0 | 0 |
| IR | 5 | 24 | 19 | 0 | 0 |
| IR | 6 | 3 | 4 | 0 | 0 |
| IR | 7 | ~200 | ~200 | 0 | 0 |
| IR | 8 | 72 | 70 | 0 | 0 |
| IR | 9 | ~ 200 | ~ 200 | 0 | 0 |
| IR | 10 | ~200 | 0 | 0 | 0 |
| IR | 11 | ~260 | ~260 | 0 | 0 |
| IR | 12 | 68 | 65 | 0 | 0 |
| KS | 1 | 1 | 1 | 0 | 0 |
| KS | 2 | 0 | 0 | 0 | 0 |
| KS | 3 | 2 | 2 | 0 | 0 |
| KS | 4 | NA | 4 | NA | 0 |
| KS | 5 | 2 | 2 | 0 | 0 |
| KS | 6 | 1 | 1 | 0 | 0 |
| KS | 7 | 17 | 16 | 0 | 0 |
| KS | 8 | 15 | 13 | 0 | 0 |
| KS | 9 | 3 | 3 | 0 | 0 |
| KS | 10 | TNTC | TNTC | 0 | 0 |
| KS | 11 | 1 | 1 | 0 | 0 |
| KS | 12 | 0 | 0 | 0 | 0 |
| KH | 1 | 0 | 0 | 0 | 0 |
| KH | 2 | 0 | 0 | 0 | 0 |
| KH | 3 | 0 | 0 | 0 | 0 |
| KH | 4 | 0 | 0 | 0 | 0 |
| KH | 5 | 0 | 0 | 1 | 1 |
| KH | 6 | 0 | 0 | 0 | 0 |
| KH | 7 | 3 | 4 | 1 | 1 |
| KH | 8 | 0 | 0 | 0 | 0 |
| KH | 9 | 0 | 0 | 0 | 0 |
| KH | 10 | 1 | 1 | 0 | 0 |
| KH | 11 | 0 | 0 | 0 | 0 |
| KH | 12 | 0 | 0 | 0 | 0 |

**Supplemental Table 8 continued**

| | | Total Colony Counts | | | |
|---|---|---|---|---|---|
| Dog | TP | Blood Agar 24 hours | Blood Agar 48 hours | MaConkey Agar 24 hours | MaConkey Agar 48 hours |
| LM | 1 | 0 | 0 | 0 | 0 |
| LM | 2 | 0 | 0 | 0 | 0 |
| LM | 3 | 1 | 1 | 0 | 0 |
| LM | 4 | 0 | 0 | 0 | 0 |
| LM | 5 | 0 | 0 | 0 | 0 |
| LM | 6 | 0 | 0 | 0 | 0 |
| LM | 7 | 0 | 0 | 0 | 0 |
| LM | 8 | 0 | 0 | 0 | 0 |
| LM | 9 | 0 | 0 | 0 | 0 |
| LM | 10 | 0 | 0 | 0 | 0 |
| LM | 11 | 0 | 0 | 0 | 0 |
| LM | 12 | 4 | 4 | 0 | 0 |
| MS | 1 | 3 | 3 | 1 | 1 |
| MS | 2 | 0 | 0 | 0 | 0 |
| MS | 3 | 3 | 4 | 0 | 0 |
| MS | 4 | 0 | 0 | 0 | 0 |
| MS | 5 | 0 | 0 | 0 | 0 |
| MS | 6 | 1 | 6 | 0 | 0 |
| MS | 7 | 0 | 0 | 0 | 0 |
| MS | 8 | 0 | 1 | 0 | 0 |
| MS | 9 | 7 | 7 | 0 | 0 |
| MS | 10 | 0 | 0 | 0 | 0 |
| MS | 11 | 0 | 0 | 0 | 0 |
| MS | 12 | 0 | 0 | 0 | 0 |
| OB | 1 | 100 | 100 | 71 | 63 |
| OB | 2 | ~315 | ~320 | ~200 | ~350 |
| OB | 3 | 25 | 25 | 11 | 11 |
| OB | 4 | 90 | 90 | 26 | 25 |
| OB | 5 | 53 | 58 | 19 | 19 |
| OB | 6 | 96 | 90 | 47 | 45 |
| OB | 7 | 58 | 57 | 36 | 36 |
| OB | 8 | 71 | 64 | 55 | 50 |
| OB | 9 | 74 | 73 | 5 | 5 |
| OB | 10 | 48 | 45 | 25 | 24 |
| OB | 11 | 34 | 36 | 8 | 8 |
| OB | 12 | 44 | 44 | 6 | 6 |
| SJ | 1 | 0 | 0 | 0 | 0 |
| SJ | 2 | 0 | 1 | 0 | 0 |
| SJ | 3 | 3 | 4 | 2 | 3 |
| SJ | 4 | 0 | 0 | 0 | 0 |
| SJ | 5 | 0 | 1 | 1 | 0 |
| SJ | 6 | 1 | 2 | 0 | 0 |
| SJ | 7 | 0 | 0 | 2 | 3 |
| SJ | 8 | 0 | 0 | 0 | 0 |
| SJ | 9 | 0 | 1 | 0 | 0 |
| SJ | 10 | 0 | 0 | 0 | 0 |
| SJ | 11 | 0 | 1 | 0 | 0 |
| SJ | 12 | 0 | 0 | 0 | 0 |

**Supplemental Table 9: Eight most common bacterial tax cultured in multiple dogs over time.** (+) taxa present; (-) taxa absent

| | Dog ID | *Streptococcus canis* | *Staphylococcus pseudintermedius* | *Curtobacterium flaccumfaciens* | *Pantoea agglomerans* | *Haemophilus haemoglobinophilus* | *Escherichia coli* | *Lysinibacillus fusiformes* | *Staphylococcus intermedius* |
|---|---|---|---|---|---|---|---|---|---|
| Females | KH | - | - | - | - | - | - | - | - |
| | EH | + | - | - | - | - | - | - | - |
| | SJ | - | - | + | + | - | - | - | - |
| | HF | - | + | - | - | - | + | + | + |
| | IO | + | - | + | + | - | - | - | - |
| | LM | + | + | - | - | + | - | - | - |
| | MS | - | + | + | - | - | - | - | - |
| Males | GC | - | - | - | - | - | - | - | - |
| | IR | + | - | - | - | - | - | + | - |
| | FC | + | - | - | - | - | + | - | - |
| | OB | + | - | - | + | + | - | - | - |
| | AbB | + | + | - | - | - | - | - | - |
| | ArB | - | - | - | - | - | - | - | - |
| | KS | + | + | - | - | - | - | - | + |

**Supplemental Table 10: Antimicrobial resistance profiles.** Resistance profiles of 48 bacterial isolates (27 species) cultured from the urine of 13 healthy dogs over 14 timepoints. NA = We did not attempt to grow isolate on this media. Isolate grew (+) or did not grow (-) on media containing the specified antimicrobial. Media contained antimicrobials at breakpoint concentrations reported in dogs and/or specific to the canine urinary tract: nalidixic acid (32 ug/mL), ciprofloxacin (4 ug/mL), amoxicillin (8 ug/mL), oxacillin (0.5 ug/mL).

| Isolate ID | Dog | Timepoint | Bacterial Species | Gram Status | Nalidixic Acid | Ciprofloxacin | Oxacillin | Amoxicillin |
|---|---|---|---|---|---|---|---|---|
| 1 | HF | TP9 | *Escherichia coli* | Gram Negative | - | - | NA | + |
| 2 | HF | TP10 | *Escherichia coli* | Gram Negative | - | - | NA | + |
| 3 | HF | TP11 | *Escherichia coli* | Gram Negative | - | - | NA | + |
| 4 | HF | TP11 | *Escherichia coli* | Gram Negative | - | - | NA | + |
| 5 | IO | TP9 | *Pantoea agglomerans* | Gram Negative | - | - | NA | + |
| 6 | KH | TP5 | *Pseudomonas aeruginosa* | Gram Negative | + | - | NA | + |
| 7 | KH | TP7 | *Pseudomonas aeruginosa* | Gram Negative | + | - | NA | + |
| 8 | KS | TP11 | *Frederiksenia canicola* | Gram Negative | - | - | NA | + |
| 9 | MS | TP1 | *Pantoea ananatis* | Gram Negative | - | - | NA | + |
| 10 | MS | TP6 | *Pseudomonas fulva* | Gram Negative | + | - | NA | + |
| 11 | OB | TP1 | *Citrobacter braakii* | Gram Negative | - | - | NA | + |
| 12 | OB | TP1 | *Citrobacter freundii* | Gram Negative | - | - | NA | + |
| 13 | OB | TP4 | *Haemophilus haemoglobinopohilus* | Gram Negative | - | - | NA | + |
| 14 | OB | TP8 | *Pantoea agglomerans* | Gram Negative | - | - | NA | + |
| 15 | OB | TP12 | *Proteus hauseri* | Gram Negative | - | - | NA | + |
| 16 | OB | TP12 | *Providencia alcalifaciens* | Gram Negative | - | - | NA | + |
| 17 | SJ | TP6 | *Pantoea agglomerans* | Gram Negative | - | - | NA | + |
| 18 | AbB | TP5 | *Staphylococcus pseudintermedius* | Gram Positive | NA | NA | - | + |
| 19 | AbB | TP12 | *Staphylococcus pseudintermedius* | Gram Positive | NA | NA | - | - |
| 20 | AbB | TP8 | *Streptococcus canis* | Gram Positive | NA | NA | - | + |
| 21 | EH | TP7 | *Bacillus megaterium* | Gram Positive | NA | NA | - | - |
| 22 | EH | TP10 | *Streptococcus canis* | Gram Positive | NA | NA | - | - |
| 23 | FC | TP1 | *Kocuria rhizophila* | Gram Positive | NA | NA | - | + |
| 24 | FC | TP11 | *Streptococcus canis* | Gram Positive | NA | NA | + | - |
| 25 | GC | TP3 | *Bacillus oceanisediminis* | Gram Positive | NA | NA | + | + |
| 26 | HF | TP6 | *Arcanobacterium canis* | Gram Positive | NA | NA | - | - |
| 27 | HF | TP8 | *Lysinibacillus fusiformis* | Gram Positive | NA | NA | + | + |
| 28 | HF | TP1 | *Staphylococcus pseudintermedius* | Gram Positive | NA | NA | - | + |
| 29 | HF | TP12 | *Staphylococcus pseudintermedius* | Gram Positive | NA | NA | - | + |
| 30 | IO | TP8 | *Curtobacterium flaccumfaciens* | Gram Positive | NA | NA | + | - |
| 31 | IO | TP3 | *Streptococcus canis* | Gram Positive | NA | NA | + | + |
| 32 | IR | TP6 | *Lysinbacillus fusiformis* | Gram Positive | NA | NA | + | + |
| 33 | IR | TP3 | *Rothia nasimurium* | Gram Positive | NA | NA | - | - |
| 34 | IR | TP4 | *Streptococcus canis* | Gram Positive | NA | NA | + | - |
| 35 | KH | TP10 | *Bacillus megaterium* | Gram Positive | NA | NA | + | - |
| 36 | KS | TP8 | *Staphylococcus auricularis* | Gram Positive | NA | NA | - | + |
| 37 | KS | TP8 | *Staphylococcus intermedius* | Gram Positive | NA | NA | + | + |
| 38 | KS | TP2 | *Staphylococcus pseudintermedius* | Gram Positive | NA | NA | + | - |
| 39 | KS | TP9 | *Staphylococcus pseudintermedius* | Gram Positive | NA | NA | + | - |
| 40 | KS | TP1 | *Streptococcus canis* | Gram Positive | NA | NA | - | - |
| 41 | LM | TP12 | *Staphylococcus pseudintermedius* | Gram Positive | NA | NA | - | - |
| 42 | LM | TP12 | *Streptococcus canis* | Gram Positive | NA | NA | - | + |
| 43 | MS | TP6 | *Curtobacterium flaccumfaciens* | Gram Positive | NA | NA | NA | - |
| 44 | MS | TP3 | *Micrococcus luteus* | Gram Positive | NA | NA | + | + |
| 45 | MS | TP1 | *Stapylococcus pseudintermedius* | Gram Positive | NA | NA | + | + |
| 46 | MS | TP9 | *Stapylococcus pseudintermedius* | Gram Positive | NA | NA | - | + |
| 47 | OB | TP7 | *Staphylococcus epidermidis* | Gram Positive | NA | NA | + | + |
| 48 | SJ | TP7 | *Curtobacterium flaccumfaciens* | Gram Positive | NA | NA | - | + |

**Supplemental Table 11: Study Follow-up.** All dogs enrolled in this study remained healthy with no urinary tract issues reported in the 18-24 months following enrollement. Follow up was not available for EH. Three dogs reported brief limited episodes of gastrointestinal (GI) distress.

| Dog ID | Any urinary tract issues observed in the past 18-24 months? | Other health issues observered in the past 18-24 months? | Date of Owner Response |
|---|---|---|---|
| **KH** | No | No | 2/16/23 (in-person) |
| **IR** | No | No | 2/16/23 (email) |
| **FC** | No | Stress related diarrhea | 2/16/23 (email) |
| **OB** | No | No | 2/17/23 (email) |
| **ArB** | No | No | 2/23/23 (in-person) |
| **AbB** | No | No | 2/23/23 (in-person) |
| **EH** | N/A | N/A | N/A |
| **MS** | No | No | 2/18/23 (email) |
| **GC** | No | No | 2/17/23 (email) |
| **IO** | No | No | 2/16/23 (email) |
| **KS** | No | Vomiting (suspected GI virus) | 2/16/23 (email) |
| **LM** | No | No | 2/16/23 (email) |
| **SJ** | No | Vomiting due to plant ingestion | 2/16/23 (text) |
| **HF** | No | No | 2/16/23 (text) |

### Supplemental Figures

**Supplemental Figure 1: Urine pH and protein correlations.** There was no significant correlation between **a)** pH and the total protein band number (*R* = 0.057, *p* = 0.36) or between **b)** pH and Tamm Horsfall concentration (*R* = -0.0099, *p* = 0.86)**,** or between **c)** pH and albumin concentration (*R* = 0.067, *p* = 0.22).

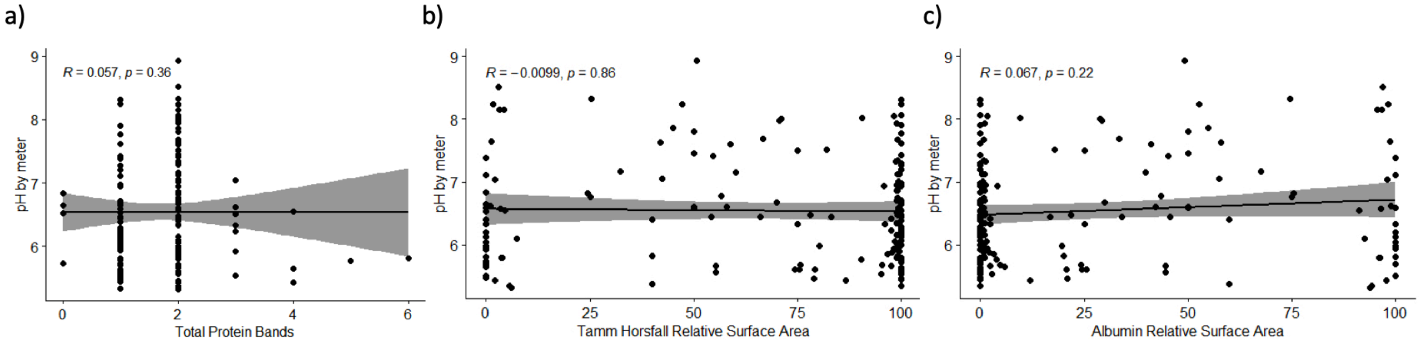
